# RNA viral communities are structured by host plant phylogeny in oak and conifer leaves

**DOI:** 10.1101/2021.12.17.473209

**Authors:** Anneliek M. ter Horst, Jane D. Fudyma, Aurélie Bak, Min Sook Hwang, Christian Santos-Medellín, Kristian A. Stevens, David M. Rizzo, Maher Al Rwahnih, Joanne B. Emerson

## Abstract

Wild plants can suffer devastating diseases, experience asymptomatic, persistent infections, and serve as reservoirs for viruses of agricultural crops, yet we have a limited understanding of the natural plant virosphere. To access representatives of locally and globally distinct wild plants and investigate their viral diversity, we extracted and sequenced dsRNA from leaves from 16 healthy oak and conifer trees in the UC Davis Arboretum (Davis, California). From *de novo* assemblies, we recovered 389 RNA-dependent RNA polymerase (RdRp) gene sequences from 384 putative viral species, and a further 580 putative viral contigs were identified with virus prediction software followed by manual confirmation of virus annotation. Based on similarity to known viruses, most recovered viruses were predicted to infect plants or fungi, with the highest diversity and abundance observed in the *Totiviridae* and *Mitoviridae* families. Phyllosphere viral community composition differed significantly by host plant phylogeny, suggesting the potential for host-specific viromes. The phyllosphere viral community of one oak tree differed substantially from other oak viral communities and contained a greater proportion of putative mycoviral sequences, potentially due to the tree’s more advanced senescence at the time of sampling. These results suggest that oaks and conifers harbor a vast diversity of viruses with as-yet unknown roles in plant health and phyllosphere microbial ecology.

## Introduction

Trees and other wild plants can act as reservoirs of viruses that can cause disease in economically important crops (Cooper and Jones 2006; Ma et al. 2019; Schoelz and Stewart 2018), but most research on plant viruses has been focused on viruses that cause disease in crops and ornamental plants (Roossinck 2014; Roossinck et al. 2015). Very little is known about the prevalence and effects of viral infection in wild plants (Prendeville et al. 2012; Roossinck 2014), even though these viruses can play important ecological roles in the phytobiome, even in asymptomatic plants (Cooper and Jones 2006; Ma et al. 2019; Schoelz and Stewart 2018). In particular, forest trees, such as oaks and conifers, have a broad ecological distribution and substantial economic importance, yet there is a relative lack of information about their associated viral diversity (Büttner et al. 2013; Aldrich and Cavender-Bares 2011; Farjon 2018; Richardson et al. 2007).

Forests are among the world’s most important ecosystems. They cover 30% of the Earth’s land surface, preserve most of Earth’s terrestrial biodiversity, are an important carbon sink, and play a role in climate regulation (Morin et al. 2018; Trumbore et al. 2015; Wingfield et al. 2015). Moreover, the forestry industry in the United States provides four percent of the total manufacturing Gross Domestic Product (GDP), which equals an estimated contribution of over $200 billion per year (Oswalt 2021). The increase in global trade has accelerated the spread of invasive pathogens to forests (Gauthier et al. 2015; Langor et al. 2014). These pathogens are either introduced by accident, and/or adapt to new host trees (Wingfield et al. 2015) and are responsible for major economic and ecological damages (Gonthier and Nicolotti 2013; Nery 2020).

With the emergence of next generation sequencing (NGS), great advances have been made in the discovery of plant viruses, and these studies have shown diverse viruses in wild plants (Ma et al. 2019; Roossinck 2012; Cooper and Jones 2006). Currently little is known about the functions of these viruses, but it is clear that viruses can be symbiotic members of their plant host microbial community (Roossinck 2011). Viruses associated with plants, including viruses that infect the plant and viruses that infect members of the plant microbiome, are known to play various roles with respect to plant health. Even though most plant viruses have been studied in the context of disease (Roossinck 2011), some plant-associated viruses have been shown to have positive, mutualistic, or neutral interactions with the plant host (Lefeuvre et al. 2019; Roossinck 2015a, 2014, 2015b), either directly (Xu et al. 2008; Westwood et al. 2013) or indirectly (Nuss 2000, 2005; Wang et al. 2019; Balogh et al. 2010; Buttimer et al. 2017). For example, the mycovirus *Cryphonectria hypovirus 1* causes hypovirulence (reduced virulence) of the fungus, *Cryphonectria parasitica*, that causes chestnut blight. (Nuss 2000). As part of the plant microbiome, bacteria and fungi can have diverse ecological interactions with the plant host, but the role of viruses that infect these plant-associated microbiota is relatively poorly understood.

Here, we used a double-stranded RNA enrichment protocol to obtain RNA viral nucleic acids from plant leaves, because most RNA viruses have a dsRNA life stage (Roossinck 2015b), including most known plant-infecting viruses and mycoviruses (Gergerich and Dolja 2006; Rubio et al. 2020). This dsRNA enrichment protocol has fostered in-depth analyses of virus-specific sequences from plants in the past (Roossinck 2012, 2014), and we chose it over other common techniques for viral community analyses with the expectation that it could facilitate better access to the phyllosphere viral community. In previous studies, enrichment of virus-like particles followed by nucleic acid extraction and sequencing has had variable results in plants (Roossinck 2017; Melcher et al. 2008), and extraction of total DNA or RNA followed by bioinformatic mining of viral sequences has resulted in less viral ‘signal’ in the sequencing data, as the majority of the sequencing reads tend to be derived from cellular organisms (Roossinck et al. 2015).

In this study, we sequenced dsRNA derived from leaves of 16 oak and conifer species in order to reconstruct RNA viral contigs and assess the RNA viral diversity within and among these host tree species. The assembled contigs were examined for RNA-dependent RNA polymerase (RdRp) genes, a conserved gene found in RNA viruses that lack a DNA stage (Starr et al. 2019), along with other viral signatures, and viral communities were compared across tree species.

## Results and discussion

### Dataset overview and viral contig recovery

To investigate the diversity and abundance of RNA viruses in conifer and oak trees, we extracted dsRNA from the leaves of 16 tree species (5 *Cupressaceae*, 5 *Pinaceae*, and 6 *Fagaceae*, commonly called cypresses, pines, and oaks, respectively, Supplementary table 1) from the UC Davis Arboretum. Samples were sequenced to an approximate depth of 6.6 Gbp each. Reads were assembled into contigs ≥ 200 bp, which were searched for viral signatures using 1) established Hidden Markov Models (HMMs) to identify viral RdRp genes (Starr et al. 2019), 2) VIBRANT (Kieft et al. 2020) virus prediction software, and 3) BLASTp and BLASTn (Altschul et al. 1990) searches against the NCBI nr (BLASTp) and nt (BLASTn) databases to manually investigate the putative viral contigs found via the first two approaches (Supplementary Figure 1). Of 186,591 total contigs ≥ 200 bp in the dataset, 1,166 were tentatively predicted as viral using approaches 1 and 2, and 202 of those were removed after manual investigation revealed an ambiguous or likely non-viral origin using approach 3, most often due to evidence that the contig was derived from a retrotransposon, not a virus. We note that, despite the dsRNA extraction that should have yielded predominantly viral RNA, many contigs were not predicted to be viral. Although we do not know the reason for certain, our virus detection approach was meant to be conservative to limit false positives, so we have probably missed some viruses, and a BLASTp search of all 192,336 predicted protein sequences revealed 69.5% of the sequences to be of likely plant origin (Supplementary Figure 1).

We detected 389 RdRp gene sequences on 384 contigs and a further 614 putative viral proteins on 580 contigs via VIBRANT, for a total of 964 putative viral contigs that passed the manual curation step. In order to maximize recovery of viral sequences that may not have assembled into contigs and to calculate the relative abundance of each putative virus in each sample, we mapped reads from each of the 16 samples to both the 964 viral contigs recovered *de novo* and 4,495 RefSeq viral genomes (Pruitt et al. 2007). We detected 963 viral contigs through read mapping (889 from the Arboretum contigs and 74 from RefSeq). The 389 RdRp sequences from the Arboretum were translated to protein and used for phylogenetic analyses, and the remaining analyses of viral populations and communities within and among trees considered the 963 viral contigs recovered through read mapping.

### Viral RdRp diversity in leaves from 16 oak and conifer species

To investigate viral diversity within the 16 tree species (Supplementary Table 1), we explored the phylogenies of the recovered RdRps in the context of known RdRps in the RefSeq database. We first translated the 389 predicted RdRp contigs into protein sequences and then dereplicated them at 99% amino acid identity (AAI) into 337 putative RdRp protein sequences, which were phylogenetically grouped with 14 viral families via BLASTp searches against RefSeq RdRp sequences. A phylogenetic tree of our RdRps and 635 RefSeq RdRps from these 14 viral families was then constructed (Figure 1A). Most of our RdRps grouped phylogenetically with unclassified viruses (n=92), followed by *Totiviridae* (n=38), *Bunyaviridae* (n=33), and *Secoviridae* (n=30) (Figure 1A). The *Totiviridae* are known to infect fungi and protozoa (Okamoto et al. 2016), the *Bunyaviridae* infect plants, insects, and vertebrates (Payne 2017), and the *Secoviridae* infect plants (Thompson et al. 2014), suggesting that both plant-infecting viruses and viruses that infect other members of the phytobiome were recovered.

**Figure 1:**
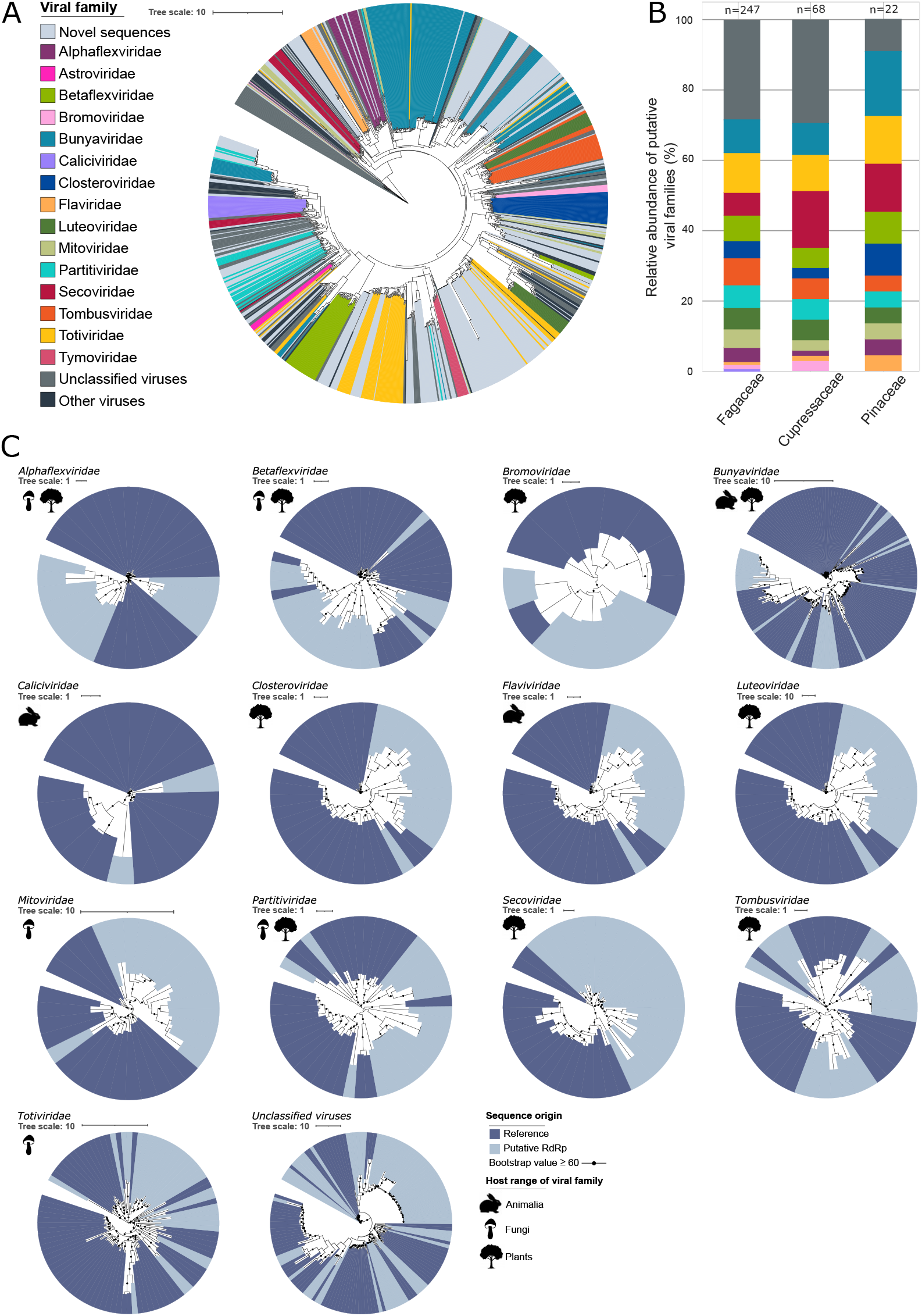
Phylogenetic classification of viral contigs, based on alignment of RdRp gene sequences from RefSeq and this dataset. A) Unrooted phylogenetic tree (concatenated predicted protein alignment) of RdRp sequences from all newly identified contigs (‘Novel sequences’, this dataset, n=337) and RefSeq (n=635). The tree is colored by viral family phylogeny from RefSeq. ‘Unclassified viruses’ and ‘Other viruses’ are also from RefSeq but did not have a taxonomic assignment or were assigned to other viral groups, respectively. B) Average relative abundances of viral taxa within tree families, based on putative taxonomic assignments for RdRp contig sequences (derived from significant BLAST hits for the RdRp gene to known RdRp sequences in RefSeq). Colors correspond to the legend in panel A. ‘Unclassified viruses’ indicate RdRps from our dataset that had significant best BLAST hits to unclassified viruses in RefSeq. Relative abundances of each RdRp-containing contig in each sample were derived from read mapping to the 384 RdRp-containing contigs, and relative abundances were summed for each taxon and averaged across tree species within each tree family to generate the stacked bar charts. Numbers at the tops of bars indicate the total number of RdRp-containing contigs detected for each tree family. C) Unrooted phylogenetic trees of RdRp protein sequences. The most significant RdRp BLAST bit score was used to assign each contig to a viral family, and phylogenetic trees were constructed separately for each family. Bootstrap support values ≥ 60% are shown as circles on nodes, and were calculated using an approximate likelihood ratio test (aLRT) with the Shimodaira–Hasegawa-like procedure (SH-aLRT), using 1000 bootstrap replicates.

In a comparison of viral taxonomic diversity according to tree family (the *Pinaceae* and *Cupressaceae* families of conifers and the *Fagaceae* family of oaks), viral taxonomic composition was similar overall, but substantially more viruses were identified from oaks than from the two conifer tree families (Figure 1B). For both the *Cupressaceae* and *Fagaceae*, the ‘taxonomic’ category with the most RdRps was the ‘unclassified viruses’ group, at 29% and 28% of total RdRps, respectively. The largest taxonomic group associated with the *Pinaceae* was the *Bunyaviridae* family (18%). Besides unclassified viruses, all three tree families had most RdRps associated with *Totiviridae, Bunyaviridae*, and *Secoviridae*, consistent with the dominance of these groups in the dataset overall.

We next sought to consider the potential hosts of the recovered viruses, based on phylogenetic affiliation of their RdRps with those of viruses with known hosts in RefSeq (Figure 1C). Of the 337 RdRps in the dataset, 32% were phylogenetically affiliated with viruses thus far only known to infect plants (*Bromoviridae, Closteroviridae, Luteoviridae, Secoviridae*, or *Tombusviridae* (Fermin 2018)), 20% were associated with viruses exclusively known to infect fungi (*Mitoviridae* or *Totiviridae* (Fermin 2018)), and 19% were aligned with viral families known to infect both plants and fungi (*Alphaflexiviridae, Betaflexiviridae, Partitiviridae* (Fermin 2018)). A further 18% grouped with unclassified viruses and thus could not be assigned to a putative host, while 10% were associated with viral families known to infect both plants and animals such as insects (*Bunyaviridae* (Payne 2017)). Overall, the dataset was dominated by putative plant and fungal viruses.

However, 2% of the RdRps were associated with viruses from the *Caliciviridae* or *Flaviviridae* (Fermin 2018) that are thus far only known to infect *Animalia* (Figure 1C). Four of these RdRps were initially aligned with the *Flaviviridae*, which are known to infect vertebrates and are transmitted by arthropods (MacLachlan 2017), and two aligned with *Caliciviridae*, which are known to only infect vertebrates (Payne 2017). Since both the *Caliciviridae* and *Flaviviridae* contain important human and animal pathogens (Desselberger 2019; MacLachlan 2017), these RdRps are well represented in the RefSeq database (Pruitt et al. 2007), and we suspected that the RdRps in our dataset could have been erroneously assigned to these groups, partly on account of this database bias. To further investigate whether these RdRps could represent true *Caliciviridae* or *Flaviviridae*, we performed a manual, web-based BLASTp search against the NCBI non-redundant protein database. Two of our RdRp sequences had significant alignments with RdRps of totiviruses, one with partitiviruses, one with mitoviruses, one with tombus-like viruses, and the last one with tymoviruses. All of these virus groups include known plant and/or fungal viruses (Martelli et al. 2002; Fermin 2018), thus all of the RdRps originally assigned to *Caliciviridae* or *Flaviviridae* were more likely to be derived from plant or fungal viruses.

Overall, results are consistent with plant and/or fungal viruses dominating the RNA viral communities in these oak and conifer phytobiomes. Interestingly, RdRps from bacteriophages were not detected in our dataset, despite potential host bacteria presumably representing a large component of the phyllosphere and endosphere microbiome (Starr et al. 2019; Chaudhry et al. 2020). We infer that either bacteriophages with RNA genomes were not abundant in these samples or that they were not amenable to the laboratory and/or bioinformatic approaches used for viral recovery. Most known bacteriophages have dsDNA genomes (Dion et al. 2020), which would not have been recovered through our dsRNA library preparation, so the lack of detection of bacteriophage RdRps does not necessarily mean that bacteriophages were not present.

### Comparing viral populations detected within and across tree families

We next sought to investigate the extent to which specific viruses were shared within and among the three tree families. We used the 389 RdRp-containing contigs, 580 putative viral contigs identified by VIBRANT (Kieft et al. 2020), and 4,495 viral genomes from RefSeq as a reference database for read mapping from the 16 dsRNA metagenomes to assess the presence of each virus in each tree. Viruses were most commonly shared among tree species within the same family, with the *Pinaceae* having the most shared viruses in the dataset (164), followed by the *Fagaceae* (78), and then the *Cupressaceae* (70) (Figure 2). After similarities within families, the *Cupressaceae* and *Pinaceae* together shared the most viral contigs (63). Both of these tree families belong to the order *Pinales* (conifers) and are more closely related to each other than they are to the *Fagaceae*. More viral contigs were shared across all three tree families (*i.e*., detected in at least three trees, with at least one species from each tree family) than were shared between the *Fagaceae* and either the *Cupressaceae* or *Pinaceae* alone. Together, these results could indicate that there are both specific, host-associated viromes that align with tree phylogeny (for example, due to the presence of specific plant metabolites and/or host-associated microbiomes that could in turn select for specific viruses) and a core virome common across tree families, perhaps due to their shared location within the UC Davis Arboretum.

**Figure 2:**
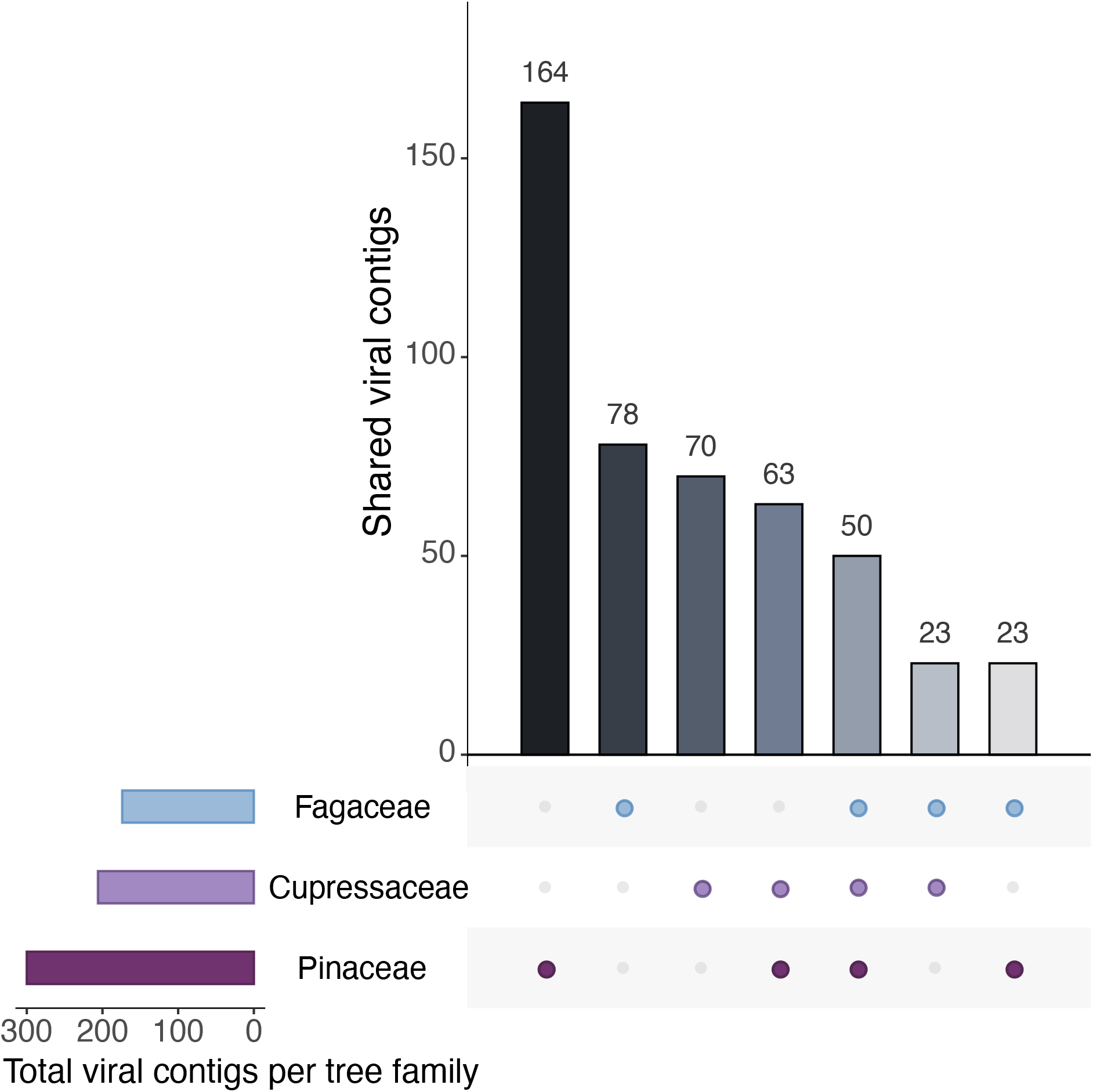
Upset plot of shared viral contig sequences between tree families based on the presence/absence version of the table of viral contig sequences in each sample (Supplementary Table 3). Colored dots below the bars indicate the tree family or families included in each bar. Only viral contigs detected in two or more tree species were included in this analysis (n=471).

### Comparing viral community composition to host tree phylogeny

Given that more viruses were shared within than among tree families, we wanted to see whether leaf viral communities would separate according to host tree phylogeny. Based on read mapping from each sample to all of the viral contigs and RefSeq viral genomes, we computed pairwise correlations between viral communities using the Pearson method and compared the resulting hierarchical cluster of viral community composition with a phylogenetic tree of the tree species, derived from the chloroplast rbcL gene (commonly used to define tree phylogeny (Manhart 1994; Kang et al. 2017)). As in the phylogenetic tree of the trees, the hierarchical cluster of viral communities showed clear separation between the oaks and the conifers, along with separation according to the two families within the conifers (*Cupressaceae* and *Pinaceae*), yielding three primary clusters of viral community composition according to tree family (Figure 3A). In fact, for all 10 conifer species, the dendrogram of viral community composition and the phylogenetic tree of trees aligned exactly, suggesting strong ties between host plant phylogeny and viral community composition, presumably due to host specificity for the viruses to the plants themselves and/or to their specific microbiomes. In contrast, within the oak (*Fagaceae*) family, the dendrogram of viral communities and the host phylogenetic tree had slight differences. We attribute these differences at least in part to differences in the phylogenetic dispersion captured in the oaks compared to the conifers: all of the oaks were from the same genus (*Quercus*), whereas the conifers spanned seven genera. Consistent with this, the three conifer species in the *Pinus* genus had viral communities that were more similar to each other than to the other two members of the *Pinaceae* family from different genera. We speculate that, at the shorter phylogenetic distances captured within a tree genus, the influence of host phylogeny on properties that would influence viral community composition was much smaller than at longer phylogenetic distances, where, in all cases, viral community composition aligned with tree phylogeny.

**Figure 3:**
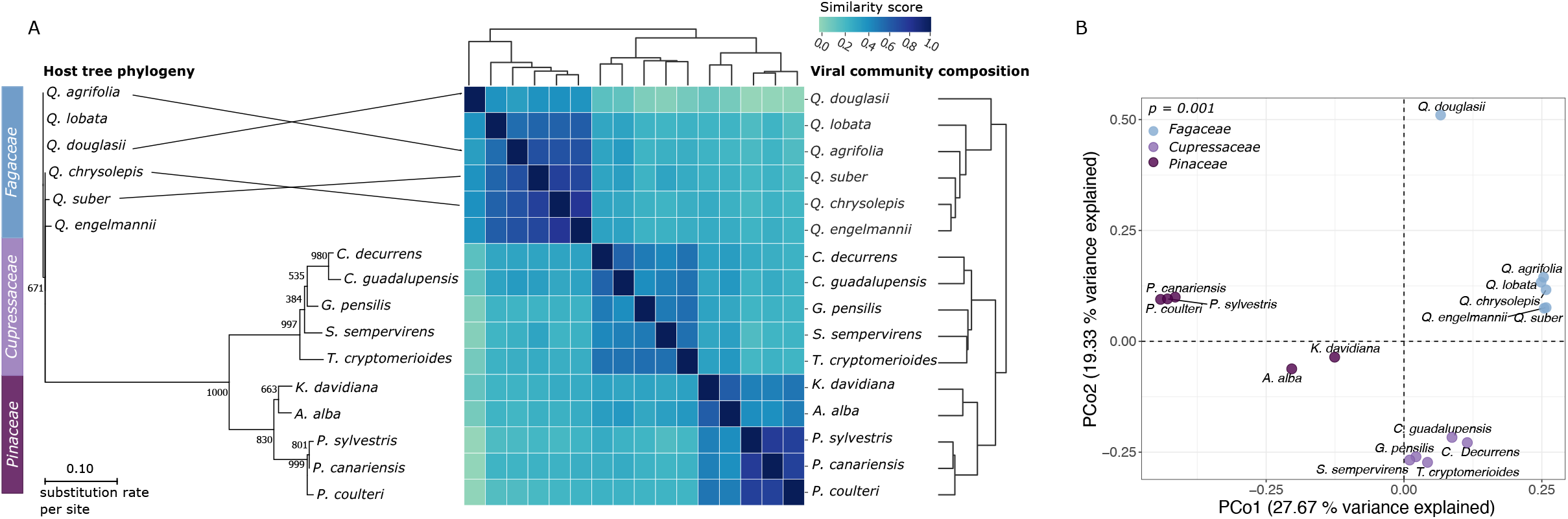
Viral community composition by tree taxonomy. A) Unrooted phylogenetic tree of tree species, based on sequence alignment of the RbcL gene (left), connected via a tanglegram (lines after tree species names; no line indicates the same row, equivalent to a horizontal straight line) to a heatmap and associated dendrograms (right and top, same dendrogram repeated) of pairwise viral community similarity between tree leaf samples. Viral community similarity was measured as Pearson similarity between pairs of samples, starting from a presence-absence matrix of 964 viral contigs in each sample, with detection patterns based on read mapping to the viral contigs. Crossed lines in the tanglegram indicate discrepancies between tree phylogeny and the viral community composition dendrogram. B) Principal Coordinates Analysis (PCoA) of viral community composition, based on pairwise Bray-Curtis dissimilarities derived from per-sample read mapping to 964 viral contigs. Each point represents a sample from one of the 16 tree species, labeled by species and colored by tree family. The p-value is from a PERMANOVA test for differences in viral community composition according to the three tree families.

To further investigate the viral communities across the 16 tree species, we performed a Principal Coordinate Analysis (PCoA, Figure 3B). This analysis now considers the relative abundances of all of the viruses recovered in the dataset through read mapping, whereas the analysis above was based solely on detection (presence/absence). Consistent with the presence/absence analysis, viral communities differed significantly by tree family (PERMANOVA p = 0.001) (Figure 3B). Viral community similarity in the PCoA plot followed approximately the same patterns as in the dendrogram derived from viral presence/absence data. However, the ‘outlier’ viral community from the *Q. douglasii* oak was particularly pronounced as separate from the other oaks in the PCoA plot, and this ‘outlier’ sample warranted further consideration, as described in the next section.

### Evidence for differences in leaf viral community composition between more senescent and less senescent trees

The viral community of the oak tree *Q. douglasii* differed substantially from the viral communities of all other oak trees (Figure 3A, 3B), likely in part because the largest number of viral contigs (333) was recovered from that tree. The second-largest number of viral contigs recovered from a tree was 251 (in *Pinus sylvestris*), and the average number for all other tree species was 130(Supplementary Figure 2). *Q. douglasii* was the only species in this study known to be drought deciduous, meaning that leaves are shed in response to drought (McCreary 2021; Abrams 1990). Consistent with the drought deciduous lifestyle resulting in earlier senescence and loss of leaves, we found that the leaves of *Q. douglasii* were in a further state of senescence compared to other samples, as shown by their brown color (Supplementary Figure 2). As plants senescence and turn into dead organic matter, saprobic fungi become more active in order to decompose this organic matter (Lindahl and Tunlid 2015), and we hypothesized that viruses of these fungi might have become more active as well.

To investigate whether the senescing *Q. douglasii* sample contained more mycoviral sequences than other samples, we focused on the subset of RdRp HMM searches with matches to *Mitoviridae*, which was the most abundant viral family in the dataset exclusively known to infect fungi. A total of 83 matches to the *Mitoviridae* RdRp HMM was found in the dataset overall, and 87% of these (72 out of 83) were recovered through read mapping in the *Q. douglasii* sample, compared to a maximum of 15% in all other samples. Of all of the *Q. douglasii* RdRp HMM matches that were recovered through read mapping (n=231), 31% matched the *Mitoviridae* HMM (72 out of 231), compared to a maximum of 16% (27 out of 170) in all other samples. Thus, both the diversity and proportion of *Mitoviridae* sequences was higher within *Q. douglasii* than in any of the other trees. We infer that *Q. douglasii* had more mycoviral sequences than any of the other trees, and we suspect that this may have been due to increased fungal and mycovirus activity within the leaves as the tree went into senescence.

### Conclusions

By extracting dsRNA from tree leaves and mining the assembled contigs for viral signatures, we recovered 964 putative viral contigs from asymptomatic oak and conifer plants. The phylogenetic affiliation of many of these viruses with known plant and fungal viruses suggests that their primary hosts are plants or fungi, potentially indicating persistent infection of the host plant and/or infection of members of the plant microbiome. Although some viruses were detected in all three tree families examined, suggesting the potential for a core virome (*e.g*., due to close proximity within the UC Davis Arboretum), most viruses that were detected in more than one tree were limited to tree species within the same tree family, suggesting host- and/or host microbiome-specific factors that could be structuring these viral communities. Interestingly, more putative mycoviruses were recovered from senescing oak leaves than from any other sample in the dataset, suggesting the potential for increased mycoviral activity coinciding with increased activity of saprobic fungi during senescence. Much viral diversity remains to be discovered, and here we have provided a framework to further investigate viral diversity in wild tree species for a more complete understanding of the plant holobiont and for identifying potential reservoirs of emergent plant diseases.

## Methods

### Sample Collection

Leaves were collected from 16 visibly healthy trees, each from a distinct species (six *Fagaceae*, five *Pinaceae*, five *Cupressaceae* trees), within the UC Davis Arboretum in Davis, CA, USA (N38° 31’ 48.3594”, W-121° 45’ 39.603” Supplementary figure 3, Supplementary table 1, Supplementary figure 4). For each plant, four to five leaves (oak leaves or conifer needles) were collected from the middle of the branches, avoiding new growth. Leaves were collected at an approximate height of 1.5 m from the ground. Mature leaves were gripped at the petiole with sterile gloved hands and pulled away from the tree without touching the exposed ends of the petiole. Leaves were immediately placed in sterile plastic bags on ice and maintained at −80°C until sample processing. Samples were processed within 96 hours of collection.

### Double-stranded RNA (dsRNA) extraction

Double-stranded RNA (dsRNA) was extracted from all samples, which had been stored at −80°C, following the protocol described by Kesanakurti et al. (Kesanakurti et al. 2016), excluding DNase and RNase treatments. Whole leaves were ground, including both the leaf surface and endosphere. RNA extracts were further prepared using a KingFisher Flex System with the MagMAX™ Plant RNA Isolation Kit (ThermoFisher Scientific, Sunnyvale, CA, USA).

### Library construction and sequencing

Per sample, a total of 700 ng per 10 μL of extracted RNA was used for cDNA library construction using the TruSeq Stranded Total RNA with Ribo-Zero Plant kit (Illumina, San Diego, CA, USA). cDNA libraries were end-repaired, adapter-ligated by unique dual-indexes, and PCR enriched. Libraries were sequenced on an Illumina NextSeq 500 platform using paired-end 150 bp format.

### Bioinformatics

Sequencing reads were quality-trimmed using Trimmomatic v0.39 (Bolger et al. 2014), using standard settings. PhiX and adapter sequences were removed using the bbduk.sh command from the bbmap package (version 38.82) (Bushnell 2014), using the k=31 and hdist=1 settings. All of the resulting clean reads, paired and unpaired, were then assembled into contigs larger than 200 bp using MEGAHIT v 1.2.9 with standard settings (Li et al. 2015). Prodigal v 2.6.3 was then used to predict genes and proteins on the assembled contigs, using the -p meta and -q flags and otherwise standard settings (Hyatt et al. 2010). Hmmer v3.2.3 (Eddy 2011) was used to perform an HMMsearch for RdRp sequences in these predicted proteins, following Starr et al., 2019 (Starr et al. 2019), with an e-value threshold of 0.00001. An HMM profile was constructed from the following RdRp PFAM numbers: Mononeg_RNA_pol [PF00946], RdRP_5 [PF07925], Flavi_NS5 [PF00972], Bunya_RdRp [PF04196], Mitovir_RNA_pol [PF05919], RdRP_1 [PF00680], RdRP_2 [PF00978], RdRP_3 [PF00998], RdRP_4 [PF02123], Viral_RdRp_C [PF17501], and Birna_RdRp [PF04197]. The recovered RdRp sequences were then clustered using USEARCH v 8.1.1861 (Edgar 2010) at 99% amino acid identity, with the centroids flag and otherwise standard settings.

In order to identify additional putative viral sequences, VIBRANT (Kieft et al. 2020) was run on all assembled contigs. VIBRANT uses searches against various databases (KEGG (Kanehisa and Goto 2000), PFAM (Finn et al. 2014), and VOG (VOGDB, release 94, vogdb.org) to identify virus-like proteins. The source code of VIBRANT was adapted so that it would consider contigs >100 bp with at least one open reading frame (ORF). BLASTp and BLASTn v2.7.1 (Altschul et al. 1990) searches were then performed using standard settings on all contigs that had an RdRp identified via HMMer, or a viral hit in the VIBRANT output, or both, using the complete NCBI nucleotide (nt), or complete protein (nr) database. All output files (HMMer, VIBRANT, BLASTn, BLASTp) were then concatenated with a custom Python script, available on github (Supplementary Table 2). These concatenated output files were then manually curated to remove contigs that had either only hypothetical proteins or matches to retroviral proteins, because retroviral matches are often retrotransposons (Starr et al. 2019).

After manually curating the viral contigs, we concatenated this file with all non-retroviral RNA viruses from RefSeq (version 203 (Pruitt et al. 2007)) and dereplicated the viral sequences using dRep (Olm et al. 2017), using the --S_algorithm ANImf -sa 0.95 -nc 0.85 settings. We then mapped clean sequencing reads to the curated, dereplicated viral contigs, using bowtie2 (Langmead and Salzberg 2012) with the --sensitive setting. A coverage table of viral relative abundances in each tree sample was then constructed using CoverM (v0.6.1, https://ecogenomic.org/m-tools), using the --min-covered-fraction 0.75 setting to ensure that at least 75% of the length of each contig was covered at least 1x by reads for detection in a given sample.

### Construction of a tree species phylogenetic tree, based on the rbcL gene

The nucleotide sequence of the rbcL gene (a chloroplast gene, used to define tree phylogeny (Manhart 1994; Kang et al. 2017)) was downloaded from the NCBI RefSeq database (version 203,(Pruitt et al. 2007)) for each of the tree species, except *Q. lobata*, for which no rbcL gene sequence was available. For *Q. lobata*, we aligned the complete chloroplast assembly (which includes the rcbL gene) with the rcbL gene sequences of the other tree species in this study, and then we manually curated and trimmed the alignment so that the sequence was in line with the other tree species sequences. MAFFT v7 (Katoh and Standley 2013) (standard settings) was used for alignment, and TrimAl v1.3 (Capella-Gutiérrez et al. 2009) with a no-gaps setting was used to remove ambiguous aligned sequences. PhyML v3.0 (Guindon et al. 2010) was used for building the tree, using bootstrap n=1000, substitution model hky85, and gamma distribution, subtree pruning, and regrafting. The tree was visualized and colored in iTol (Letunic and Bork 2019).

### Construction of phylogenetic trees based on the viral RdRp gene

All putative RdRp sequences (n=348) were aligned with all viral RdRp protein sequences from RefSeq v 203 (Pruitt et al. 2007), using MAFFT v7 (Katoh and Standley 2013) with the G-INS-i algorithm, as in (Shi et al. 2016). All alignments were trimmed using TrimAl v1.3 (Capella-Gutiérrez et al. 2009). These trimmed alignments only contained the RdRp and its neighboring conserved domains, and no ambiguously aligned regions. The best-fit model of amino acid substitution was determined for each sequence alignment using ProtTest 3.4 (Abascal et al. 2005). Phylogenetic trees were then inferred, using the maximum likelihood approach (ML) with FastTree v 2.3 (Price et al. 2009), and an approximate likelihood ratio test (aLRT) with the Shimodaira–Hasegawa-like procedure was used for branch support. Trees were visualized and colored in iTol (Letunic and Bork 2019).

For the viral family phylogenetic trees, all putative RdRp sequences underwent a BLASTp search against RdRp sequences from RefSeq v 203 (Pruitt et al. 2007), and the best BLAST hit for each sequence was retained. A BLAST hit against a known RdRp sequence was found for all but 10 putative RdRp sequences. The 10 putative RdRp sequences without a BLAST hit were included in the alignment with ‘Unclassified viruses’. When two or more BLAST hits received an identical bitscore, BLAST hits were manually inspected to verify that both hits came from the same viral family, and if not, we retained the hit with the highest percent identity. Separate alignments were then constructed for each viral family, with each of our sequences aligned with RdRps from the family that had the highest BLAST hit e-value. Each of the family-level trees was constructed, using the same phylogenetic tree-building methods described above.

### Statistical analyses and figure generation

The following statistical analyses were performed in R using the vegan (Oksanen et al. 2016) package: viral sequence coverage tables were standardized using the decostand function with the Hellinger method, Bray-Curtis dissimilarities were calculated using the vegdist function, and the dissimilarity matrix was used for Principal Coordinates Analysis of viral community composition. Permutational multivariate analyses of variance (PERMANOVA) were performed with the adonis function. Figures were generated in R, using the ggplot2 package (Wickham 2016), except for the heatmap figure, which was created using Python, using the seaborn, scipy, and pandas libraries (code available on GitHub).

The presence-absence matrix was generated using the coverage table and converting any value above zero to one for presence and keeping the value as zero for absence. The presence-absence data was used to create the Upset plot (Figure 2) using the UpsetR library (Conway et al. 2017). Pairwise correlations between viral communities were calculated from the presence-absence matrix using the Pearson method from the seaborn (https://seaborn.pydata.org) library, and a dendrogram corresponding to the heatmap was created using the clustermap function of the seaborn library.

## Data accessibility

Raw sequencing reads are available on NCBI under BioProject number PRJNA771411, and sequence processing and statistical analysis code can be found on GitHub (https://github.com/AnneliektH/Arboretum).

## Acknowledgements

We thank Bryce Falk and Richard Bostock for contributing to discussions while this project was being developed. This work was primarily funded by the UC Davis College of Agricultural and Environmental Sciences and Department of Plant Pathology (new lab start-up to JBE) and was partially supported by USDA NIFA Hatch project number CA-D-PPA-2464-H. Partial support was also provided by USDA NIFA grant number 2021-67013-34815-0 and U.S. Department of Energy, Office of Science, Office of Biological and Environmental Research grant numbers DE-SC0021198 and DE-SC0020163 (grants to JBE).

## Supplementary figures

**Supplementary figure 1: Schematic diagram of the data processing pipeline**

**Supplementary figure 2: Number of viral contigs recovered and number of clean sequencing reads per sample.**

**Supplementary figure 3: Images of leaf samples within 24 hours of sample collection**

**Supplementary figure 4: Sampling locations.** Sampling locations within the UC Davis

Arboretum in Davis, California, USA. Boxes indicate the sampled tree’s common name, its scientific name, the phylogenetic family to which the tree belongs, and the general location of the tree’s natural habitat.

**Supplemental table 1: Sample information and sequencing results for the 16 trees**

**Supplemental table 2: Relevant viral prediction output for each of the putative viral contigs. Viral prediction was based on RdRp hmm profiles, VIBRANT output, BlastP output and BlastN output**

**Supplemental table 3: Normalized viral abundance table based on read mapping**

**Supplemental table 4: Predicted taxonomy for each of the proteins with a HMM RdRp match, based on best BLAST hit against RefSeq RdRps**

## References

Abascal, F., Zardoya, R., and Posada, D. 2005. ProtTest: selection of best-fit models of protein evolution. Bioinformatics. 21:2104–2105.

Abrams, M. D. 1990. Adaptations and responses to drought in Quercus species of North America. Tree Physiol. 7:227–238.

Aldrich, P. R., and Cavender-Bares, J. 2011. Quercus. In Wild Crop Relatives: Genomic and Breeding Resources: Forest Trees, ed. Chittaranjan Kole. Berlin, Heidelberg: Springer Berlin Heidelberg, p. 89–129.

Altschul, S. F., Gish, W., Miller, W., Myers, E. W., and Lipman, D. J. 1990. Basic local alignment search tool. J. Mol. Biol. 215:403–410.

Balogh, B., Jones, J. B., Iriarte, F. B., and Momol, M. T. 2010. Phage therapy for plant disease control. Curr. Pharm. Biotechnol. 11:48–57.

Bolger, A. M., Lohse, M., and Usadel, B. 2014. Trimmomatic: a flexible trimmer for Illumina sequence data. Bioinformatics. 30:2114–2120.

Bushnell, B. 2014. BBMap: a fast, accurate, splice-aware aligner. Lawrence Berkeley National Lab.(LBNL), Berkeley, CA (United States). Available at: https://www.osti.gov/biblio/1241166.

Buttimer, C., McAuliffe, O., Ross, R. P., Hill, C., O’Mahony, J., and Coffey, A. 2017. Bacteriophages and Bacterial Plant Diseases. Front. Microbiol. 8:34.

Büttner, C., Bargen, S. von, Bandte, M., and Mühlbach, H. P. 2013. Forest diseases caused by viruses. In Infectious forest diseases, Wallingford: CABI, p. 50–75.

Capella-Gutiérrez, S., Silla-Martínez, J. M., and Gabaldón, T. 2009. trimAl: a tool for automated alignment trimming in large-scale phylogenetic analyses. Bioinformatics. 25:1972–1973.

Chaudhry, V., Runge, P., Sengupta, P., Doehlemann, G., Parker, J. E., and Kemen, E. 2020. Shaping the leaf microbiota: plant–microbe–microbe interactions. J. Exp. Bot. 72:36–56.

Conway, J. R., Lex, A., and Gehlenborg, N. 2017. UpSetR: an R package for the visualization of intersecting sets and their properties. Bioinformatics. 33:2938–2940.

Cooper, I., and Jones, R. A. C. 2006. Wild Plants and Viruses: Under-Investigated Ecosystems. In Advances in Virus Research, Academic Press, p. 1–47.

Desselberger, U. 2019. Caliciviridae Other Than Noroviruses. Viruses. 11 Available at: http://dx.doi.org/10.3390/v11030286.

Dion, M. B., Oechslin, F., and Moineau, S. 2020. Phage diversity, genomics and phylogeny. Nat. Rev. Microbiol. 18:125–138.

Eddy, S. R. 2011. Accelerated Profile HMM Searches. PLoS Comput. Biol. 7:e1002195.

Edgar, R. 2010. Usearch. Available at: https://www.osti.gov/biblio/1137186.

Farjon, A. 2018. The Kew Review: Conifers of the World. Kew Bull. 73:8.

Fermin, G. 2018. Host range, host--virus interactions, and virus transmission. Viruses. :101.

Finn, R. D., Bateman, A., Clements, J., Coggill, P., Eberhardt, R. Y., Eddy, S. R., et al. 2014. Pfam: the protein families database. Nucleic Acids Res. 42:D222–30.

Gauthier, S., Bernier, P., Kuuluvainen, T., Shvidenko, A. Z., and Schepaschenko, D. G. 2015. Boreal forest health and global change. Science. 349:819–822.

Gergerich, R. C., and Dolja, V. V. 2006. Introduction to plant viruses, the invisible foe. Plant Health Instr. Available at: https://www.apsnet.org/edcenter/disandpath/viral/introduction/Pages/PlantViruses.aspx

Gonthier, P., and Nicolotti, G. 2013. Infectious Forest Diseases. CABI.

Granoff, A., and Webster, R. G. 1999. Encyclopedia of Virology. Elsevier.

Guindon, S., Dufayard, J.-F., Lefort, V., Anisimova, M., Hordijk, W., and Gascuel, O. 2010. New algorithms and methods to estimate maximum-likelihood phylogenies: assessing the performance of PhyML 3.0. Syst. Biol. 59:307–321.

Hyatt, D., Chen, G.-L., Locascio, P. F., Land, M. L., Larimer, F. W., and Hauser, L. J. 2010. Prodigal: prokaryotic gene recognition and translation initiation site identification. BMC Bioinformatics. 11:119.

Kanehisa, M., and Goto, S. 2000. KEGG: kyoto encyclopedia of genes and genomes. Nucleic Acids Res. 28:27–30.

Kang, Y., Deng, Z., Zang, R., and Long, W. 2017. DNA barcoding analysis and phylogenetic relationships of tree species in tropical cloud forests. Sci. Rep. 7:12564.

Katoh, K., and Standley, D. M. 2013. MAFFT multiple sequence alignment software version 7: improvements in performance and usability. Mol. Biol. Evol. 30:772–780.

Kesanakurti, P., Belton, M., Saeed, H., Rast, H., Boyes, I., and Rott, M. 2016. Screening for plant viruses by next generation sequencing using a modified double strand RNA extraction protocol with an internal amplification control. J. Virol. Methods. 236:35–40.

Kieft, K., Zhou, Z., and Anantharaman, K. 2020. VIBRANT: automated recovery, annotation and curation of microbial viruses, and evaluation of viral community function from genomic sequences. Microbiome. 8:90.

Langmead, B., and Salzberg, S. L. 2012. Fast gapped-read alignment with Bowtie 2. Nat. Methods. 9:357–359.

Langor, D. W., Cameron, E. K., MacQuarrie, C. J. K., McBeath, A., McClay, A., Peter, B., et al. 2014. Non-native species in Canada’s boreal zone: diversity, impacts, and risk. Environ. Rev. 22:372–420.

Lefeuvre, P., Martin, D. P., Elena, S. F., Shepherd, D. N., Roumagnac, P., and Varsani, A. 2019. Evolution and ecology of plant viruses. Nat. Rev. Microbiol. 17:632–644.

Letunic, I., and Bork, P. 2019. Interactive Tree Of Life (iTOL) v4: recent updates and new developments. Nucleic Acids Res. 47:W256–W259.

Li, D., Liu, C.-M., Luo, R., Sadakane, K., and Lam, T.-W. 2015. MEGAHIT: an ultra-fast single-node solution for large and complex metagenomics assembly via succinct de Bruijn graph. Bioinformatics. 31:1674–1676.

Lindahl, B. D., and Tunlid, A. 2015. Ectomycorrhizal fungi - potential organic matter decomposers, yet not saprotrophs. New Phytol. 205:1443–1447.

Manhart, J. R. 1994. Phylogenetic analysis of green plant rbcL sequences. Mol. Phylogenet. Evol. 3:114–127.

Martelli, G. P., Sabanadzovic, S., Abou-Ghanem Sabanadzovic, N., Edwards, M. C., and Dreher, T. 2002. The family Tymoviridae. Arch. Virol. 147:1837–1846.

Ma, Y., Marais, A., Lefebvre, M., Theil, S., Svanella-Dumas, L., Faure, C., et al. 2019. Phytovirome Analysis of Wild Plant Populations: Comparison of Double-Stranded RNA and Virion-Associated Nucleic Acid Metagenomic Approaches. J. Virol. 94

McCreary, D. D. (2021, September 6) Native California oaks losing leaves early. University of California Available at: https://ucanr.edu/blogs/blogcore/postdetail.cfm?postnum=8276

MacLachlan, N. J., & Dubovi, E. J. (2017). Chapter 29 - Flaviviridae. Fenner’s Veterinary Virology (Fifth Edition), Elsevier, p. 525–545.

Melcher, U., Muthukumar, V., Wiley, G. B., Min, B. E., Palmer, M. W., Verchot-Lubicz, J., et al. 2008. Evidence for novel viruses by analysis of nucleic acids in virus-like particle fractions from Ambrosia psilostachya. Journal of Virological Methods. 152:49–55

Miller, W. A. 1999. LUTEOVIRUS (LUTEOVIRIDAE). In Encyclopedia of Virology (Second Edition), eds. Allan Granoff and Robert G. Webster. Oxford: Elsevier, p. 901–908.

Morin, X., Fahse, L., Jactel, H., Scherer-Lorenzen, M., García-Valdés, R., and Bugmann, H. 2018. Long-term response of forest productivity to climate change is mostly driven by change in tree species composition. Sci. Rep. 8:5627.

Nery, F. M. B. 2020. Viral diversity in tree species. Available at: https://repositorio.unb.br/handle/10482/40579.

Nuss, D. L. 2000. Hypovirulence and Chestnut Blight: From the Field to the Laboratory and Back. In Fungal Pathology, ed. J. W. Kronstad. Springer Netherlands, p. 149–170.

Nuss, D. L. 2005. Hypovirulence: mycoviruses at the fungal-plant interface. Nat. Rev. Microbiol. 3:632–642.

Okamoto, K., Miyazaki, N., Larsson, D. S. D., Kobayashi, D., Svenda, M., Mühlig, K., et al. 2016. The infectious particle of insect-borne totivirus-like Omono River virus has raised ridges and lacks fibre complexes. Sci. Rep. 6:33170.

Oksanen, J., Blanchet, F. G., Friendly, M., Kindt, R., Legendre, P., McGlinn, D., et al. 2016. vegan: Community Ecology Package. R package version 2.4-3. Vienna: R Foundation for Statistical Computing.

Olm, M. R., Brown, C. T., Brooks, B., and Banfield, J. F. 2017. dRep: a tool for fast and accurate genomic comparisons that enables improved genome recovery from metagenomes through de-replication. ISME J. 11:2864–2868.

Oswalt, S., USDA, (2021, July 19) The state of the Forest. Available at: https://www.usda.gov/media/blog/2019/04/22/state-forest.

Payne, S. 2017. Viruses: From Understanding to Investigation. Academic Press.

Prendeville, H. R., Ye, X., Morris, T. J., and Pilson, D. 2012. Virus infections in wild plant populations are both frequent and often unapparent. Am. J. Bot. 99:1033–1042.

Price, M. N., Dehal, P. S., and Arkin, A. P. 2009. FastTree: computing large minimum evolution trees with profiles instead of a distance matrix. Mol. Biol. Evol. 26:1641–1650.

Pruitt, K. D., Tatusova, T., and Maglott, D. R. 2007. NCBI reference sequences (RefSeq): a curated non-redundant sequence database of genomes, transcripts and proteins. Nucleic Acids Res. 35:D61–5.

Richardson, D. M., Rundel, P. W., Jackson, S. T., Teskey, R. O., Aronson, J., Bytnerowicz, A., et al. 2007. Human Impacts in Pine Forests: Past, Present, and Future. Annu. Rev. Ecol. Evol. Syst. 38:275–297.

Roossinck, M. J. 2017. Deep sequencing for discovery and evolutionary analysis of plant viruses. Virus Res. 239:82–86.

Roossinck, M. J. 2014. Metagenomics of plant and fungal viruses reveals an abundance of persistent lifestyles. Front. Microbiol. 5:767.

Roossinck, M. J. 2015a. Move over, bacteria! Viruses make their mark as mutualistic microbial symbionts. J. Virol. 89:6532–6535.

Roossinck, M. J. 2015b. Plants, viruses and the environment: Ecology and mutualism. Virology. 479-480:271–277.

Roossinck, M. J. 2012. Plant virus metagenomics: biodiversity and ecology. Annu. Rev. Genet. 46:359–369.

Roossinck, M. J. 2011. The good viruses: viral mutualistic symbioses. Nat. Rev. Microbiol. 9:99–108.

Roossinck, M. J., Martin, D. P., and Roumagnac, P. 2015. Plant Virus Metagenomics: Advances in Virus Discovery. Phytopathology. 105:716–727.

Rubio, L., Galipienso, L., and Ferriol, I. 2020. Detection of Plant Viruses and Disease Management: Relevance of Genetic Diversity and Evolution. Front. Plant Sci. 11:1092.

Schoelz, J. E., and Stewart, L. R. 2018. The Role of Viruses in the Phytobiome. Annual Review of Virology. 5:93:111.

Shi, M., Lin, X.-D., Tian, J.-H., Chen, L.-J., Chen, X., Li, C.-X., et al. 2016. Redefining the invertebrate RNA virosphere. Nature. 540:539–543.

Starr, E. P., Nuccio, E. E., Pett-Ridge, J., Banfield, J. F., and Firestone, M. K. 2019. Metatranscriptomic reconstruction reveals RNA viruses with the potential to shape carbon cycling in soil. Proc. Natl. Acad. Sci. U. S. A. 116:25900–25908.

Thompson, J. R., Kamath, N., and Perry, K. L. 2014. An evolutionary analysis of the Secoviridae family of viruses. PLoS One. 9:e106305.

Trumbore, S., Brando, P., and Hartmann, H. 2015. Forest health and global change. Science. 349:814–818.

Wang, X., Wei, Z., Yang, K., Wang, J., Jousset, A., Xu, Y., et al. 2019. Phage combination therapies for bacterial wilt disease in tomato. Nat. Biotechnol. 37:1513–1520.

Westwood, J. H., McCann, L., Naish, M., Dixon, H., Murphy, A. M., Stancombe, M. A., et al. 2013. A viral RNA silencing suppressor interferes with abscisic acid-mediated signalling and induces drought tolerance in Arabidopsis thaliana. Mol. Plant Pathol. 14:158–170.

Wickham, H. 2016. ggplot2: elegant graphics for data analysis. data.

Wingfield, M. J., Brockerhoff, E. G., Wingfield, B. D., and Slippers, B. 2015. Planted forest health: The need for a global strategy. Science. 349:832–836.

Xu, P., Chen, F., Mannas, J. P., Feldman, T., Sumner, L. W., and Roossinck, M. J. 2008. Virus infection improves drought tolerance. New Phytol. 180:911–921.

