## supplementary figures 1-4 for "RNA viral communities are structured by host plant phylogeny in oak and conifer leaves"

4 supplementary figures can be found in this file, 4 supplementary tables can be found in supplementary_tables.xlsx.

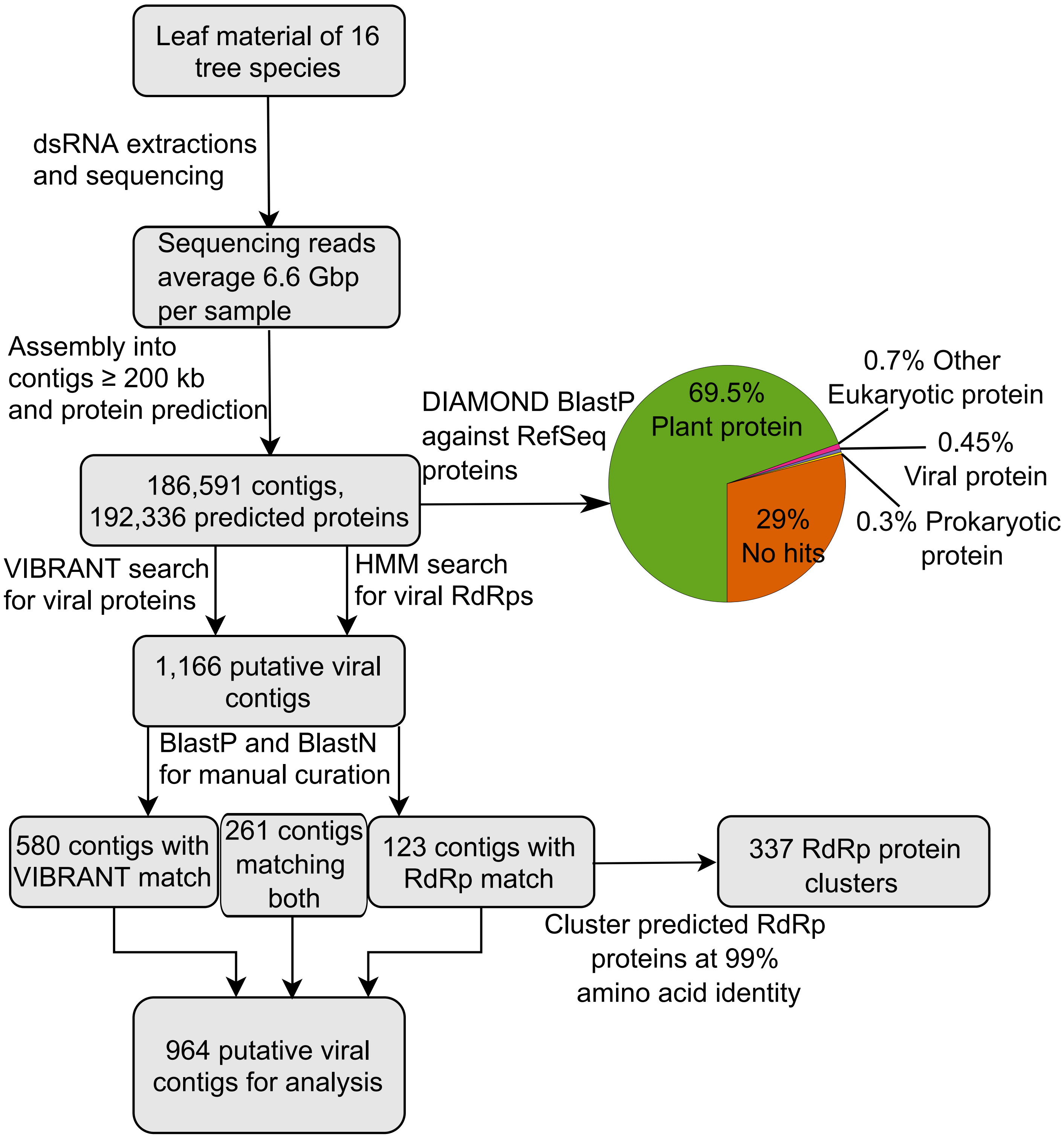


**Supplementary figure 1: Schematic diagram of the data processing pipeline**

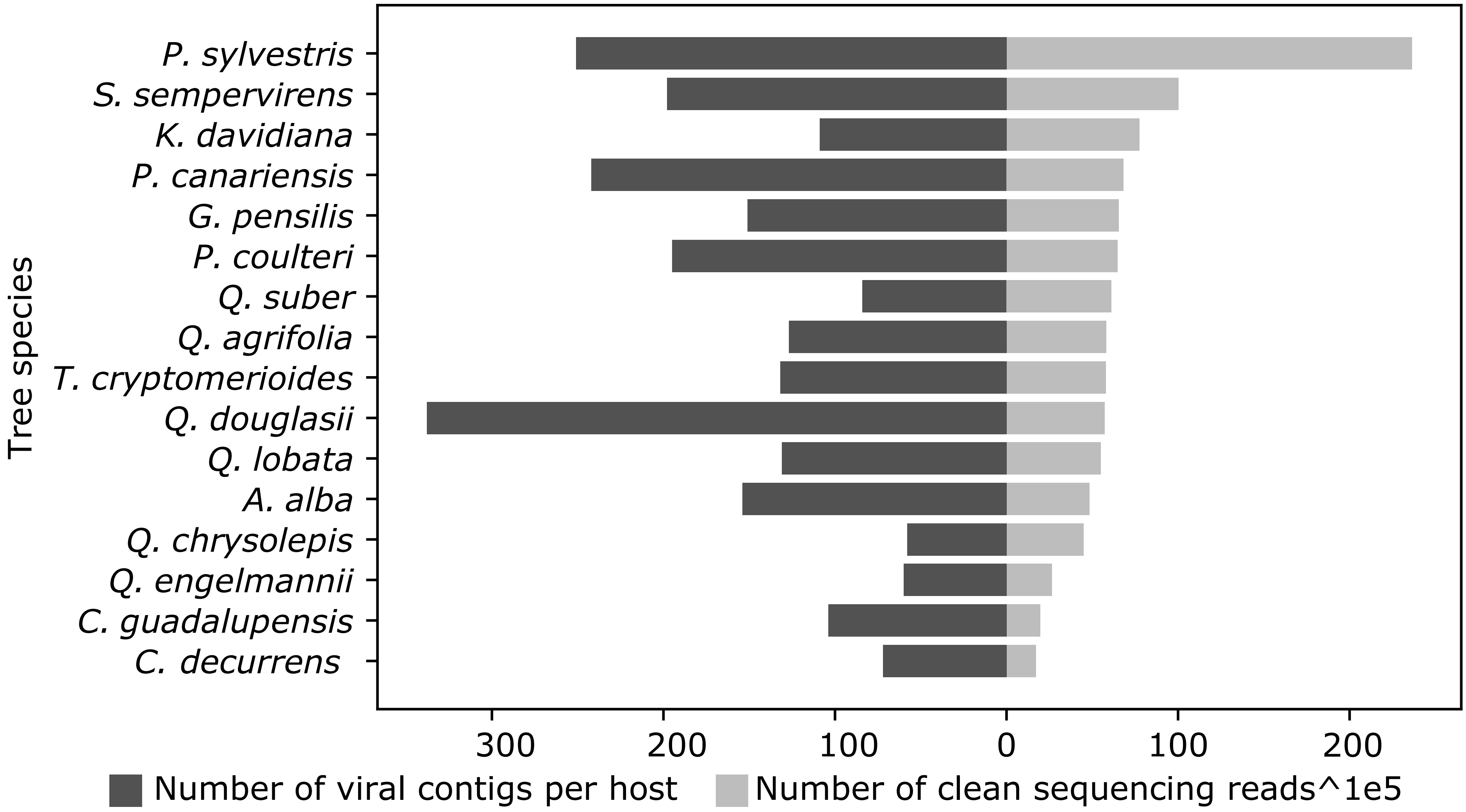


**Supplementary figure 2: Number of viral contigs recovered and number of clean sequencing reads per sample.**

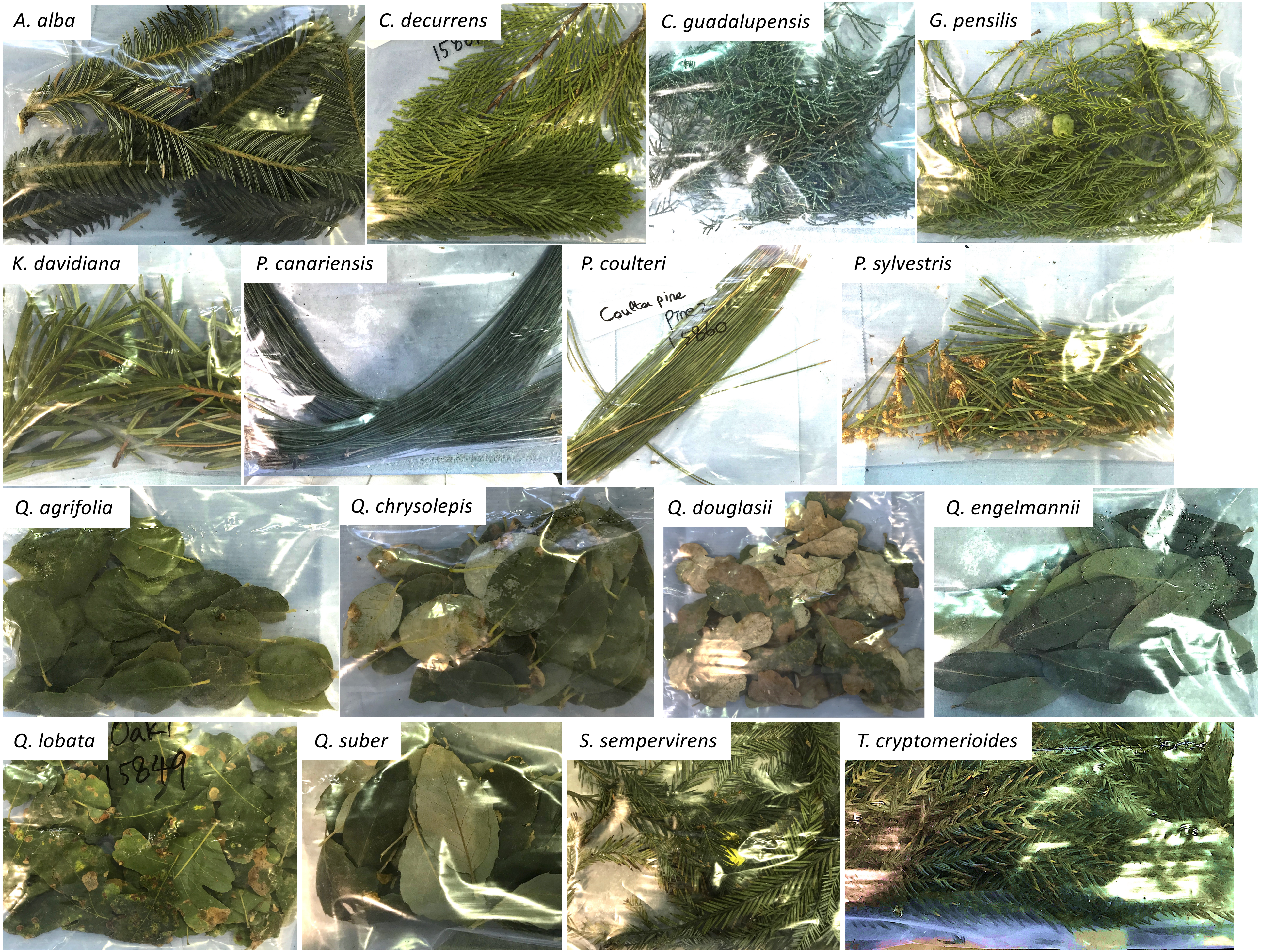


**Supplementary figure 3: Images of leaf samples within 24 hours of sample collection**

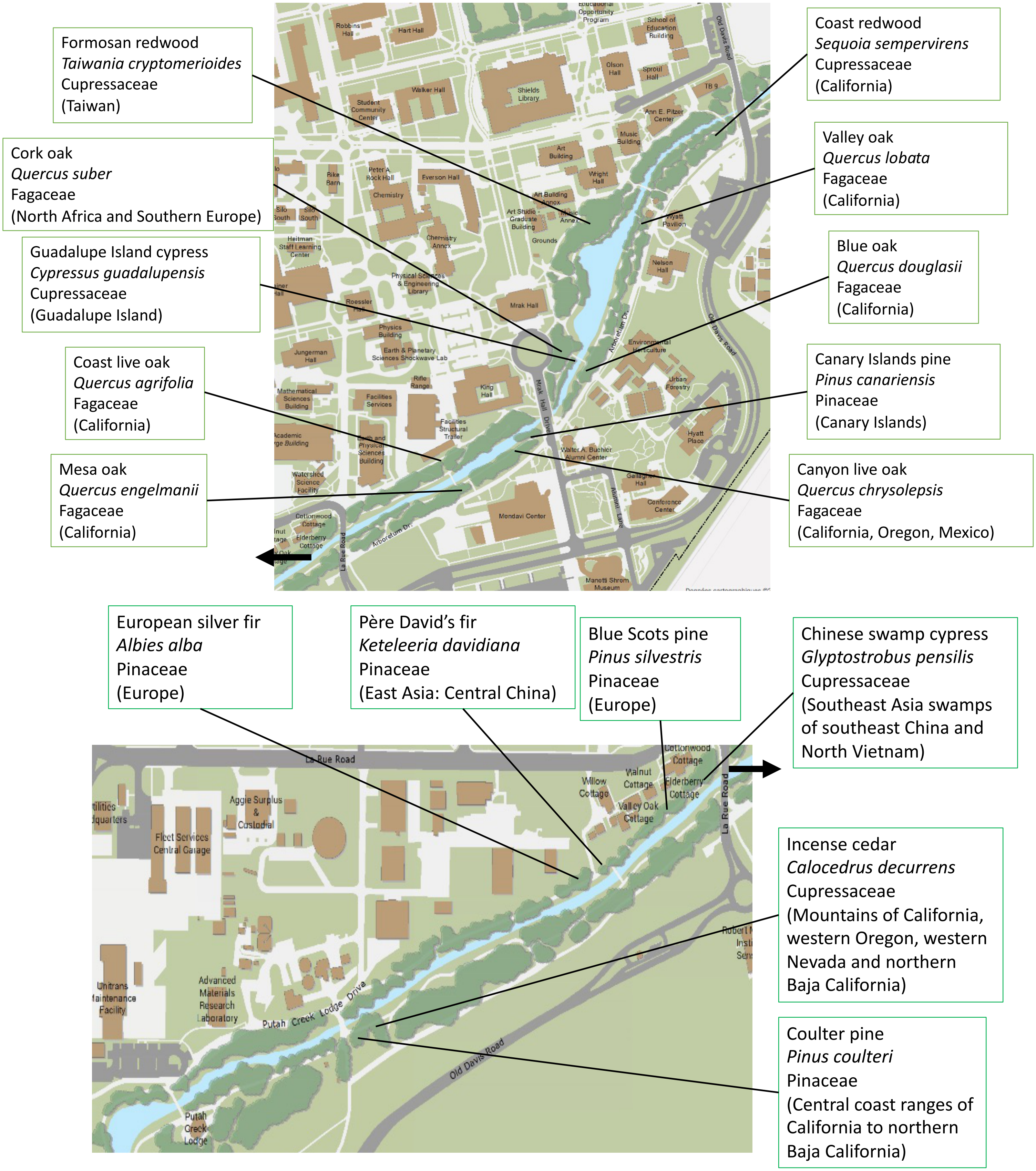


**Supplementary figure 4: Sampling locations.** Sampling locations within the UC Davis Arboretum in Davis, California, USA. Boxes indicate the sampled tree's common name, its scientific name, the phylogenetic family to which the tree belongs, and the general location of the tree's natural habitat.
